# Roughness and dynamics of proliferating cell fronts as a probe of cell-cell interactions

**DOI:** 10.1101/2020.10.26.354878

**Authors:** Guillaume Rapin, Nirvana Caballero, Iaroslav Gaponenko, Benedikt Ziegler, Audrey Rawleigh, Ermanno Moriggi, Thierry Giamarchi, Steven A. Brown, Patrycja Paruch

## Abstract

Juxtacellular interactions play an essential but still not fully understood role in both normal tissue development and tumour invasion. Using proliferating cell fronts as a model system, we explore the effects of cell-cell interactions on the geometry and dynamics of these one-dimensional biological interfaces. We observe two distinct scaling regimes of the steady state roughness of *in-vitro* propagating Rat1 fibroblast cell fronts, suggesting different hierarchies of interactions at sub-cell lengthscales and at a lengthscale of 2–10 cells. Pharmacological modulation significantly affects the proliferation speed of the cell fronts, and those modulators that promote cell mobility or division also lead to the most rapid evolution of cell front roughness. By comparing our experimental observations to numerical simulations of elastic cell fronts with purely short-range interactions, we demonstrate that the interactions at few-cell lengthscales play a key role. Our methodology provides a simple framework to measure and characterise the biological effects of such interactions, and could be useful in tumour phenotyping.

Proliferating cell fronts underpin wound healing, tumour invasion, and morphogenesis, and are governed by a complex and still not fully understood interplay of multiple mechanical and chemical factors. From a physics perspective, these cell fronts can be described as elastic interfaces in disordered media. Thus, analysing their geometric properties and dynamics can reveal the characteristic lengthscales of the dominant interactions, and point towards the underlying biological pathways. Here, we show that the roughness of proliferating fronts of Rat1 fibroblasts is governed by two different hierarchies of interactions, with distinct behaviour at sub-cell and few-cell lengthscales. Using in-vitro scratch assays, we find moreover that pharmacological modulators significantly affect the proliferation speed of the cell fronts as well as the evolution of their roughness, increased when cell-cell communication via gap junctions is perturbed, and decreased when cell division is repressed. We determine that our experimental observations cannot be reproduced by numerical simulations of elastic cell fronts with purely short-range interactions, demonstrating the key role of juxtacellular interactions at few-cell lengthscales. Our approach provides a simple framework to measure and characterise the biological effects of such interactions, and could be useful in tumour phenotyping.

## I. INTRODUCTION

The physics of elastic interfaces in disordered media provides a powerful general framework for understanding systems as diverse as eroding coastlines, propagating cracks [1], domain walls in ferroic materials [2, 3], and proliferating cell and bacterial colonies [4, 5]. In such systems, competition between the flattening effects of elasticity and the fluctuations induced by the disorder landscape leads to jerky, highly non-linear dynamics and self-affine interfacial roughening, defined by characteristic scaling exponents [6]. Formally, the geometrical fluctuations of a roughened interface are described by its monovalued transverse displacements *u*(*z*) from an elastically optimal flat configuration along the longitudinal coordinate *z*, as schematically illustrated in Fig. 1(a). The scaling properties reflected in the evolution of the correlation function *B*(*r*) of relative displacements Δ*u*(*r*) = *u*(*z*) − *u*(*z* + *r*) as a function of observation lengthscale *r* are quantified by the roughness exponent *ζ*, with

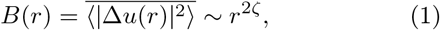

where ⟨…⟩ denotes an average over *z*, and 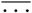 an average over disorder realizations, when appropriate.

**FIG. 1.**
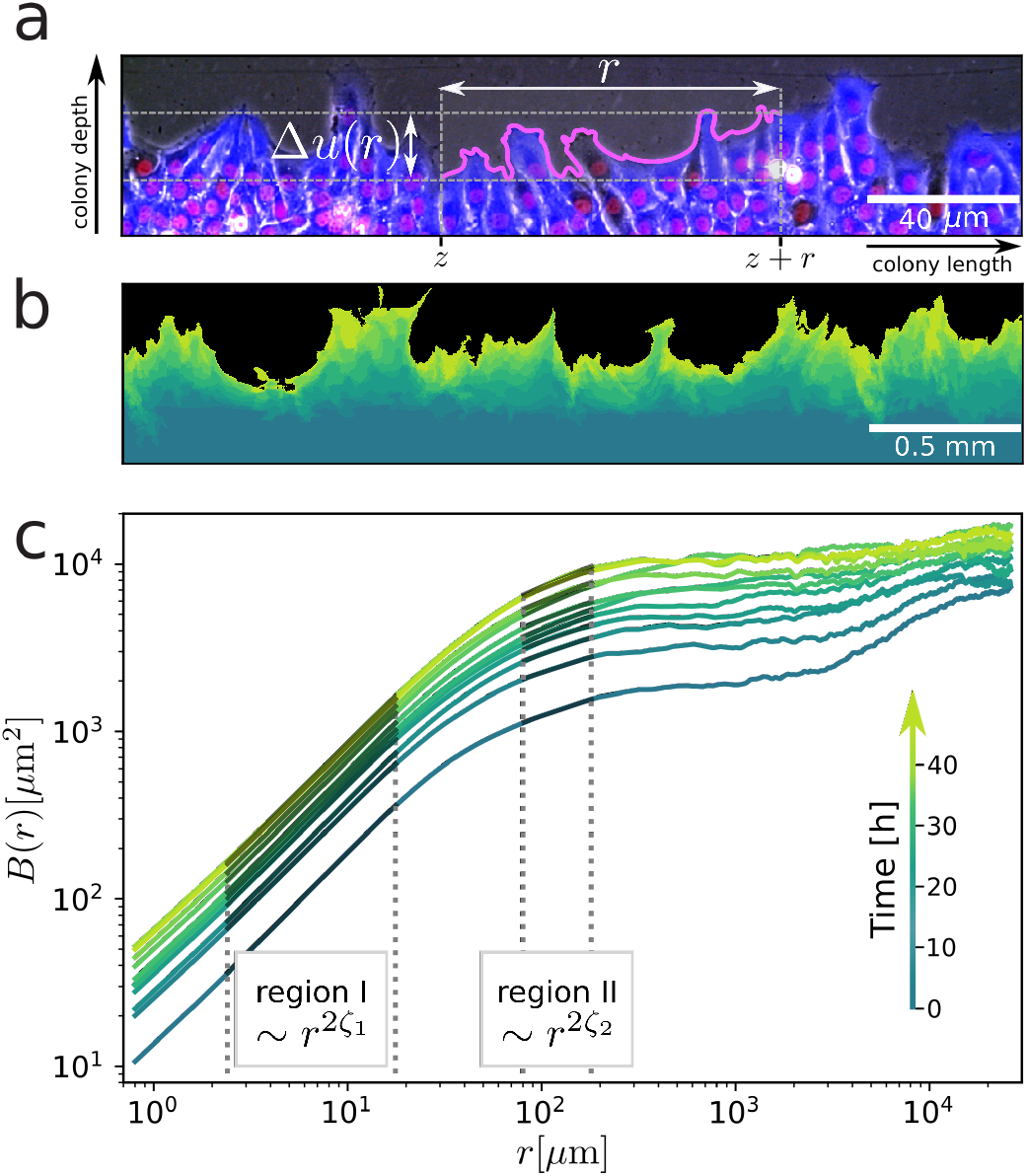
Proliferating Rat1 fibroblast cell front under control conditions. (a) Optical phase microscopy image of a section of the cell front, overlaid with cytoplasm (blue) and nuclei (red) fluorescence. The relative displacements Δ*u*(*r*) = *u*(*z*) − *u*(*z*+*r*) are measured between pairs of points separated by a distance *r*, and their correlations give a quantitative assessment of the cell front roughness. (b) Superposition of successive fluorescence microscopy images taken over 40 hours. (c) Average cell front roughness 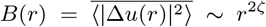 showing two distinct regions of power law scaling. In (b) and (c) the same time scale of dark to light green is used.

In physical systems, the values of the scaling exponents have been clearly related to the dimensionality of the system and the universality class of the disorder [6–8], linked to specific types of material defects [2, 9, 10]. In living systems, however, where understanding cellular proliferation and migration is a crucial first step to modelling biologically important processes such as wound healing, tumour growth, and morphogenesis, the situation is far less clear, with a great richness and complexity of interactions at both sub- and super-cell lengthscales making such assignments difficult.

At both the level of single cells and colonies, the mechanical role of the cytoskeleton and cell-cell junctions has long been recognised [11–13], forming a continuous network transferring forces between cells and through the substrates [14–16]. Growth pressure exerted by continued cell division has also been observed to drive aspects of morphogenesis [17] and colony motion [18], in some cases competing with the effects of ‘leader’ cells to give rise to highly heterogeneous proliferation of the cell front in scratch assay measurements [14, 19]. In addition, chemical signalling between cells, for example through gap junctions or juxtacrine singalling molecules [20, 21], provides further pathways for short to medium-range interactions.

Previous studies of cell front roughening have observed fractal dimension *D*_*f*_ = 1.25 ± 0.05, and a roughness exponent of 0.85 in propagating linear cell fronts and 0.5 ± 0.05 on propagating circular colonies, exhibiting a velocity of between 12 to 13.5μm/h [22, 23]. However, even with pharmacological modulation to target specific types of cellular interactions, the continued change in the shape and size distribution of the Vero cells used in these experiments, as well as their evolution from a 2D to a 3D colony, makes it very difficult to identify the effects of potentially quite different hierarchies of interactions at different lengthscales. To better explore cell front geometry at both sub-cell and super-cell lengthscales, and compare it to existing theoretical models of interfacial roughening [6], as well as numerical simulations of growing cell colonies [24, 25], a simpler model system of a purely 2D proliferating colony of homogeneous cells would be useful.

Here, we report on the roughness and dynamics of such a 2D system: propagating Rat1 fibroblast cell fronts studied in an in-vitro scratch assay over multiple orders of lengthscales (from 1 μm to 2 cm) and several days, with and without pharmacological modulation targeting cell division rate, cell motility and mechanical intracell and intercell force transmission, as well as certain types of cell-cell communication. We find two different regimes of power law scaling in the cell front roughness, with distinct values of the exponent: below the average cell size *ζ* ≈ 0.58 for all conditions, and between 5 to 10 cells, *ζ* varies from 0.13 to 0.25, with a marked effect of pharmacological modulation. At the same time, these inhibitors also have significant effects on cell front dynamics, for example increasing proliferation speed when cell-cell communication via gap junctions is perturbed, and decreasing proliferation speed when cell division is repressed.

We compare our experimental observations to numerical simulations using a vertex model developed to reproduce the elastic response of epithelial sheets [24, 25], based on the energy balance between optimal cell area, cell membrane elasticity, and cell-cell adhesion. We construct a “phase diagram” of cell front proliferation and roughness under a wide set of model parameters and demonstrate that while a single region of power law scaling with relatively high *ζ* ~ 0.74 values can be reproduced using only elasticity and nearest-neighbour interactions, the appearance of a second region with with lower *ζ* cannot. These results suggest that the geometry of propagating cell fronts is governed by two different hierarchies of interactions. Roughening at sub-cell lengthscales is consistent with a description based on cell membrane elasticity and short range interactions, but at few-cell lengthscales a decreased roughening points to the importance of collective interactions extending beyond nearest neighbours.

## II. RESULTS

### A. Experimental set-up for cell front analysis

Cell fronts were prepared from Rat1 fibroblasts, modified to express green fluorescent protein (GFP) in their cytoplasm and with their nuclei fluorescently marked using Hoecht stain H33258. Individual, initially flat fronts were created by lifting off a silicone insert from confluent plates of cells, and these fronts were imaged every four hours for more than two days using a motorised stage fluorescent microscope, with a resolution of 0.8 μm, as detailed in Methods IV. After image processing of the full scratch assay panorama of 6.0 cm (see S2), we extract the spatial position of the cell front *u*_*front*_(*z*), and approximate it by a univalued function *u*(*z*) to counter the effects of overhangs, as described in S3. Calculating the roughness function *B*(*r*) = ⟨|Δ*u*(*r*)|^2^⟩ for each front allows us to extract the roughness exponent *ζ* and its evolution with time, while the average displacement of the front ⟨*u*(*z*)⟩ with respect to the initial position allows us to extract the front dynamics. Both the roughness and dynamics are highly reproducible for a given set of conditions, as discussed in S5. Where possible, *B*(*r*) and ⟨*u*(*z*)⟩ are averaged over measurements carried out independently on different cell fronts.

### B. Cell fronts under control conditions

To establish the generic behaviour of the cell fronts, we first measured them under control conditions with no pharmacological modulation. As can be seen in Fig. 1(b), the cell fronts evolve over 40 hours from a flat initial configuration (dark green) towards an increasingly rough geometry (light green) characterized by multiple protrusions and semicircular cavities. Some cavities, like the one in the left of the image, appear to act as local pinning sites with relatively little front displacement from the initial position compared to the surrounding regions. While we do observe overhangs, most of the cell fronts present a univalued position function. We note that mitosis events, visible via the characteristic ‘bowtie’ shape of the mitotic spindle in the nuclear fluorescence channel, appear to occur at a distance of at least a few cells in from the front, but not in the cells directly at the front

The increasing front roughness is reflected in the increasing value of *B*(*r*) at all lengthscales as a function of time, as shown in Fig. 1(c), more marked in the first 12 hours, then slowing to reach an apparent steady state after about 30 hours. From a sliding analysis of an uncertainty-weighted fit of *B*(*r*) in windows of equivalent size on a logarithmic scale (as detailed in S6, we identify two distinct regions of power-law scaling. In region *I*, for sub-cell lengthscales between 2.4 μm and 18 μm, we find a roughness exponent value of *ζ*_1_ ≈ 0.58, which remains essentially constant with time (Fig. 2(c)). In region *II*, for few-cell lengthscales between 80 μm and 180 μm, we observe significantly lower values of the roughness exponent *ζ*_2_ ≈ 0.20–0.25, increasing slightly as the front proliferates (Fig. 2(d)). At high lengthscales above 300 μm, *B*(*r*) appears to saturate. While initially this saturation can be attributed to the globally flat configuration imposed by the experimental setup (with occasional artefacts related to the slight curvature of the patterning insert, as detailed in S4), once a higher value *B*(*r*) steady state is reached, saturation suggests a physical upper limit on the power law growth of cell front roughness.

Dynamically, the front shows a linear progression of displacement from its initial position, as can be seen in Fig. 3. The average colony depth is obtained by averaging the front position over its full length and comparing this to the average initial position of the front. We measure an average front velocity of 6.5 μm/h.

**FIG. 2.**
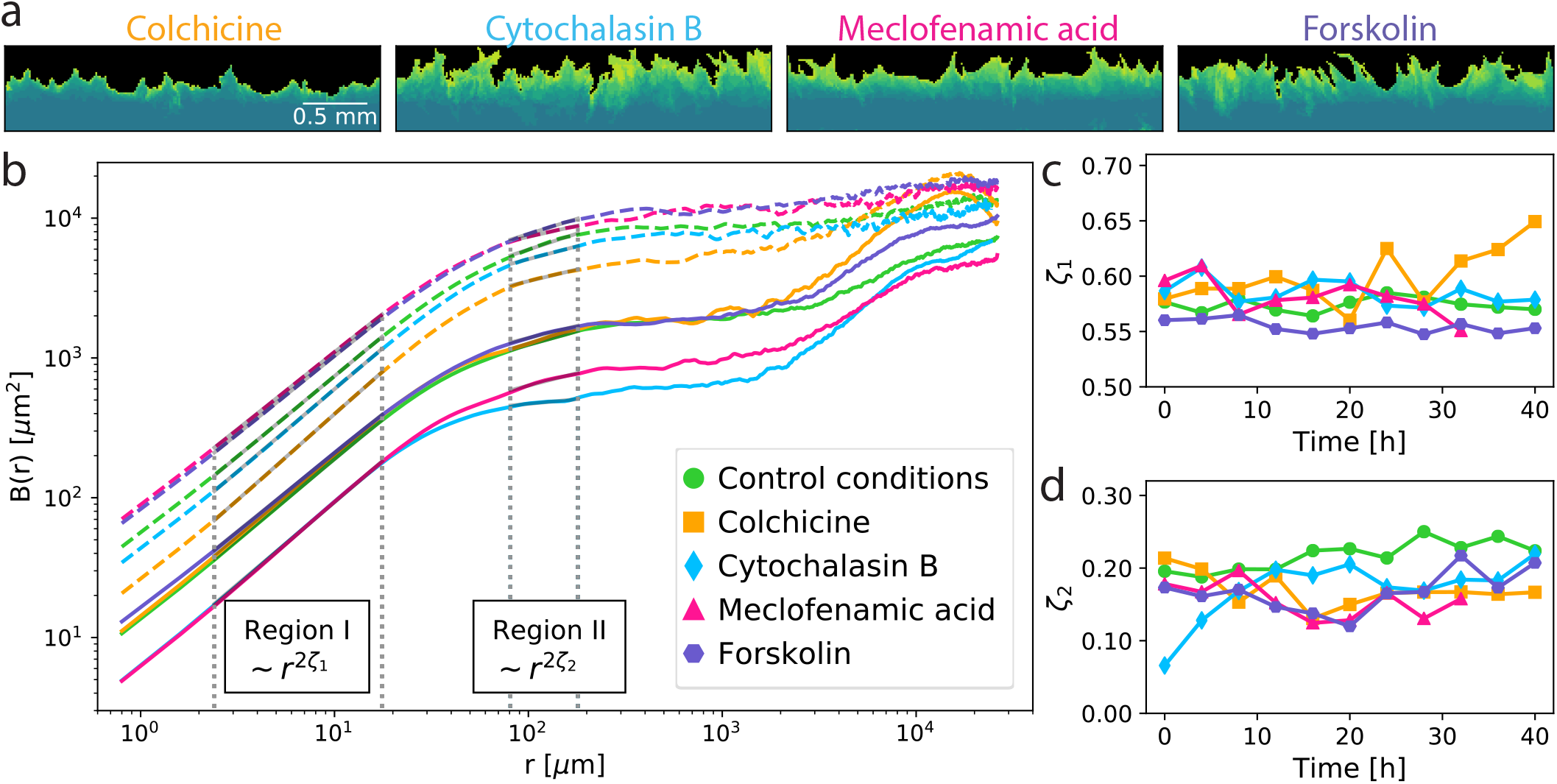
Proliferating Rat1 fibroblast cell fronts after pharmacological modulation. (a) Superposition of successive fluorescence microscopy images taken over 40 hours, with earlier front pictured in dark green and more recent ones in lighter green. (b) Cell front roughness *B*(*r*) at 0 and 24 hours for the four different inhibitors, showing again two distinct power-law scaling regions. (c,d) Roughness exponents *ζ* as a function of time for cell fronts under control conditions (green) and after exposure to cytochalasin-B (blue), colchicine (orange), meclofenamic acid (magenta), and forskolin (purple) at sub-cell lengthcales ((c), *ζ*_1_) and few-cell lengthscales ((d), *ζ*_2_).

**FIG. 3.**
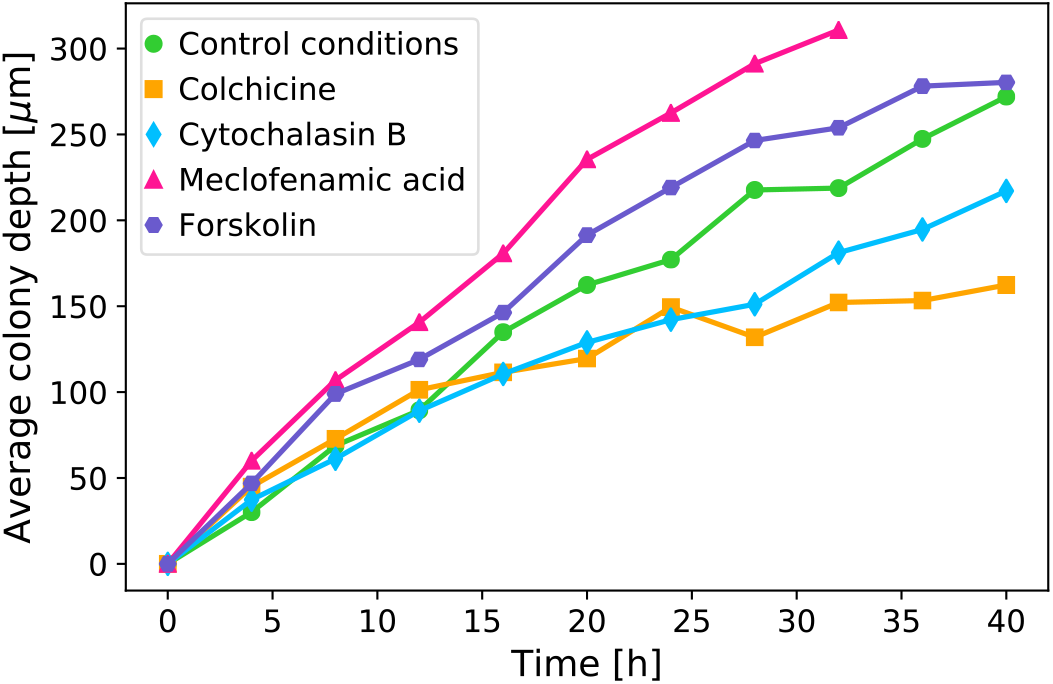
Dynamics of proliferating Rat1 epithelial cell fronts. Average colony depth as a function of time, showing the speed of proliferation is strongly affected by pharmacological modulation.

### C. Pharmacological effects on the roughness and dynamics of cell fronts

To probe the contributions of different cellular signaling pathways to cell-cell interactions at different lengthscales, we next compared the cell fronts proliferating under control conditions to cell fronts modulated using a selection of pharmacological agents. Colchicine, which binds to tubulin and thus inhibits microtubule polymerisation (in particular crucial for spindle formation during mitosis) [26], was chosen to target cell division rates, thus decreasing associated growth pressure in the cell colony. Cytochalasin B, which disrupts the polymerisation and network build-up of actin filaments [27], was chosen to inhibit both cell motility and the transmission of mechanical forces between neighbouring cells. Meclofenamic acid (MFA) inhibits gap junctions communication [28] and was chosen to disrupt this pathway of electrochemical signal transmission, previously shown to propagate logarithmically to neighboring cells up to ten neighbors distant [29]. Finally, forskolin, which activates adenyl cyclase thereby creating the signalling molecule c-AMP, an upstream activator of diverse stress and growth signalling pathways [30], was chosen in order to globally affect cell metabolism. Confluent cell cultures were incubated with each inhibitor after lift-off patterning to establish the cell front, at concentrations reported by others and verified by toxicological titration experiments in our laboratory (see S1), then imaged an analysed identically to the cell fronts under control conditions.

We observe that already with a visual inspection of the cell front morphology from fluorescence microscopy images (see Fig. 2(a), effects of the inhibitors can be discerned. Compared to control conditions, the colchicine-treated front shows overall much less proliferation, moving very slowly but coherently, with the cells appearing to stay in close contact to each other, and forming very limited protrusions. The MFA- and forskolin-treated fronts appear to proliferate more. We observe that forskolin also appears to strongly affect the shape of the cells, and the resulting front shows higher aspect ratio protrusions, present already in the initial configuration. As the front proliferates, these protrusions elongate in some parts forming few-cell-length fingers, separated by regions which appear to move very little from their initial position. The MFA-treated front, in contrast, is visually comparable to the control conditions front, although with slightly smaller protrusions. The cytochalasin Btreated front forms short spiky protrusions but at higher density, with an apparently more ‘localised’ pinning effect, and appears to move in a more incoherent fashion, laterally as well as forward, resulting in a higher number of overhangs.

Quantitatively, the inhibitor effects can be seen in the variations of the roughness *B*(*r*) for the different fronts, shown in Fig. 2(b) both at initialisation (*t* = 0 datasets, shown in solid lines) and after evolution (*t* = 24 hours datasets, shown as dashed lines). While the initial roughness is to at least some degree affected by the execution of the lift-off step (see S5 for comparisons of reproducibility between fronts), what is important here is the comparison between the roughness of the initial configuration and as the front evolves with time. Initially, the MFA- and forskolin-treated fronts appear to have the lowest roughness, but increases to the highest roughness values after 24 hours. The cytochalasin-B-treated front, initially showing the lowest *B*(*r*) values, also roughens much more significantly with time. The colchicine-treated front, in contrast, initially showing the highest *B*(*r*) values, evolves much more slowly than the others, and after 24 hours is the least rough of the fronts.

Interestingly, while the effects on the magnitude of *B*(*r*) are quite strong, overall much less variation is seen in the values of the roughness exponents. In region *I*, in particular, *ζ*_1_ values are almost identical in the initial configuration for all the fronts, varying between 0.55 and 0.6, and little change is observed as the front evolves with time, except perhaps for the colchicine-treated front, whose *ζ*_1_ exponent appears to increase to 0.65 after 40 hours of front proliferation, but also presents the highest variability. At few-cell lengthscales in region *II*, we find that fronts exposed to inhibitors generally show lower *ζ*_2_ values than fronts under control conditions. Also, while for the control condition front *ζ*_2_ evolves very slightly and approximately continuously from 0.20 to 0.25 over the 40 hour experiment duration, for the cytochalasin-B-treated front, a significant increase in *ζ*_2_ from 0.07 to 0.2 occurs in the first 10 hours, after which time the exponent value remains stable within the error bars. With MFA, colchicine and forskolin, the variations appear to be mostly within the error bars, although with potentially a small decrease of *ζ*_2_ values from 0.2 to 0.15 over the first 20 hours.

The most significant effects of the inhibitors are observed in the dynamics of the fronts, as can be seen in Fig. 3. Both the colchicine- and cytochalasin-B-treated fronts show a marked decrease in proliferation rates, down to average velocities of 3.8 μm/h and 5 μm/h, respectively. Conversely, the front treated with MFA shows strongly increased proliferation, with an average velocity of 10 μm/h, and the forskolin front a slight increase to 7.2 μm/h.

### D. Numerical simulations: the importance of cell-cell interactions

To better understand the role played by cell-cell interactions in the roughness and dynamics of the observed cell fronts, we performed numerical simulations using the vertex model implemented by the *epicell* package, chosen for its capability to reproduce the mechanical response of epithelial cell sheets [24, 25].

The Hamiltonian approach implemented in *epicell*, with

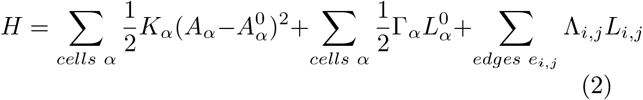

is based on energy minimisation with a balance between the cell area elasticity determined by the area elasticity coefficient *K*_*α*_, cell perimeter contractility determined by Γ_*α*_, and line tension along cell-cell junctions given by Λ_*i,j*_, further detailed in S8. The result of such an approach is shown in Fig. 4(a), with an initially regular two-dimensional hexagonal cell arrangement (i) evolving as a function of time - through cell divisions and relaxation - to a steady-state configuration (ii) with a qualitative geometry that is comparable to experimental observations of Rat1 cell fronts (iii).

**FIG. 4.**
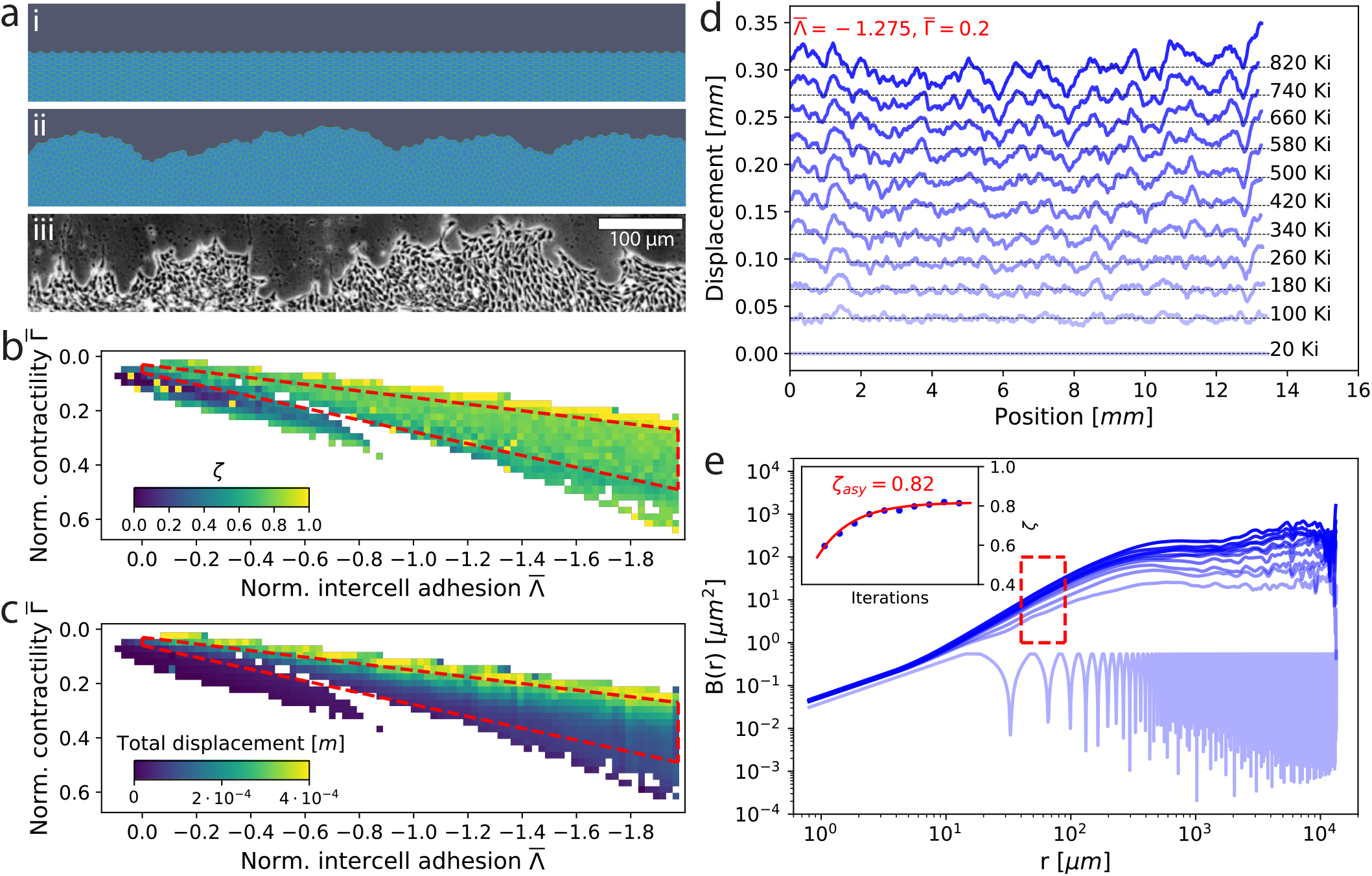
Numerical simulations of propagating cell fronts. (a) Initially flat fronts of regular two-dimensional hexagonal cells (i) were allowed to evolve under energy balance conditions [24] to a steady state rough configuration (ii), visually comparable to the Rat1 fibroblast cell fronts imaged by phase contrast (iii). Phase diagram of the (b) roughness exponent *ζ* and (c) total displacement of the cell front simulations as a function of the normalized intercell adhesion 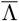 and normalized cell contractility 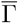. The lower branch corresponds to pathological scenarios in which little cell division occurs, accompanied by negligible front motion and roughening. The upper branch corresponds to non-pathological scenarios in which the colony grows and the front proliferates to reach an apparent steady state rough configuration and linear velocity. White space corresponds to unsuccessful simulations (see S8). The dashed red-coloured bounding box identifies the main region of stability and physical behaviour. (d) Evolution of a full ≈ 14*mm* simulated cell front for 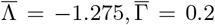 typical of the upper non-pathological branch, progressively roughening as a function of iterations from the initial flat configuration. Selected fronts are shown at intervals of 80000 iterations (80 Ki), demonstrating a linear velocity of the interface. (e) Average roughness 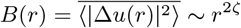 of the fronts in (d), evolving from a periodic function reflecting the initial configuration to a steady state presenting a single region of power law scaling. The evolution of the roughness exponent *ζ*, extracted by fitting the region indicated by the red box, is shown in the inset.

We chose the simulation parameters to reflect the physics of the observed system: optimal cell size was set to the average cell size of the Rat1 fibroblasts, and the cell area elasticity was taken from previous results reproducing experimentally observed mechanical strain in cell cultures [24]. As these model parameters govern govern the specific physics of the system, we explored a large range of normalized cell contractility 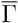 and normalized inter-cell adhesion 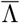 parameter combinations, yielding phase diagrams such as those shown in 4(b,c). White space corresponds to unsuccessful simulations, caused either by unphysical parameters or lack of numerical convergence. Two branches are apparent in all phase diagrams. The lower branch generally exhibits aberrant behaviours, such as limited cell division, cell sheet contraction instead of colony growth, and negligible front motion and roughening. The upper branch is generally well-behaved, and corresponds to normal behavior where the culture grows and the front proliferates, reaching a steady state roughness over time. We therefore defined an area of stability and physically relevant behaviour, bounded by the dashed red box indicated on the phase diagrams, within which we further analysed the evolution of the cell front roughness.

Each successful simulation was run for 12 hours or 20000 divisions, and then analysed individually, as illustrated in Fig. 4(d) for the full ≈ 14 mm simulated cell front with 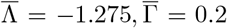, corresponding to a typical non-pathological behaviour. The front position was extracted at regular time intervals, here 80000 iterations (80 Ki), and shows a constant proliferation velocity as well as visibly steady-state roughness configuration at the final time steps, usually after 500 Ki. The average roughness *B*(*r*) was computed at each time step, starting with a highly correlated behaviour due to the initial front periodicity. Since the cells in the simulation are flat-sided polygons, roughness at sub-cell lengthscales has no biophysical relevance. As a function of time, *B*(*r*) generally tends towards a steady state behaviour with power law scaling at low lengthscales, and a transition towards a roughness saturation at around 0.5 mm.

The roughness exponent *ζ* was extracted as in the *invitro* experiment from the region highlighted by the red box shown in Fig. 4(e). Last, the evolution of *ζ* as a function of time was fitted by an exponential function to extract the asymptotic value of the roughness exponent *ζ_asy_*.

The phase diagram of asymptotic roughness exponents for each set of conditions can be seen in Fig. 4(b), and shows a large region of uniform value in the stable branch bounded by the red dashed box. As discussed above, this region corresponds to non-pathological cases, with the corresponding total displacement in Fig. 4(c) increasing both with lower contractility as well as lower inter-cell adhesion. The mean exponent within this region 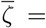 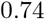, with standard deviation *σ* = 0.09.

The overall behaviour of the cell front simulations seems to suggest that power law scaling with relatively high *ζ* values can be reproduced using a simple Hamiltonian based on mechanical properties and nearest-neighbor inter-cell coupling. Although the model includes no external disorder, the simulations produce some internal disorder, and the values of *ζ* we obtain are in the same range as for example those of the so called random bond disorder universality class for which *ζ* = 2/3. However, the simulations fail to capture the experimentally observed lower roughness exponent values, and of course the presence of two distinct scaling regimes. These results imply that the geometry of *in-vitro* propagating cell fronts is governed by two different hierarchies of interactions. The experimentally observed roughening at sub-cell lengthscales can plausibly be described as a balance of cell membrane elasticity and short range interactions. However, at few-cell lengthscales, collective interactions appear to be necessary to decrease roughening and establish a second region of power law scaling with lower values of the roughness exponent.

## III. DISCUSSION

Our measurements of pharmacologically modulated cell fronts point to a complex interplay of interactions at multiple scales, and between static and dynamic parameters. Unsurprisingly, cell division appears to be a strong driver of front movement, in agreement with past observations in other systems [29]. At intermediate length scales of 5–10 cells, those modulators that promote cell mobility or division also lead to the most rapid evolution of cell front roughness. For example, inhibiting cell division with colchicine results in smaller changes of *B*(*r*) at intermediate lengthscales (5–10 cells) over time. In reverse, inhibiting cell-cell communication via gap junctions appears to both significantly promote cell front mobility and leads to the most rapid evolution of cell front roughness *B*(*r*) with time. Thus, our results also suggest that communication across these intermediate lengthscales may play an important role in determining cell front movement, and by inference biological processes such as tumor invasion, wound healing, and tissue development. We note that dye diffusion assay studies of gap junctions networks in rat kidney cells [29] have shown characteristic effective concentration lengthscales of the order of 200 μ, in the range where we observed the second power law growth region of *B*(*r*). Forskolin, activating the same cellular pathways as the most common class of G protein-coupled receptors responding to diffusible extracellular ligands, affects *B*(*r*) in the same range.

At longer lengthscales (1000 μ) where we observe the cell front flattening over time, we propose that these changes might arise from specific growth behaviors and not as simply an artefact of the initially defined flat front configuration. Indeed, while *B*(*r*) continues to increase with time, as can be seen in Fig. 1, this flattening becomes more, rather than less apparent. Overall, when cavities are present in the front, instead of proliferation globally in the direction perpendicular to the line of the front, the cells appear to actively redirect and flow towards the cavity, filling it and effectively ‘smoothing’ the front geometry at high length scales, as can be seen in S7. The specific physical or biological determinants of proliferation region and direction remains a fascinating question meriting further study.

Finally, at short lengthscales of less than two cells, from the analysis of the numerical simulations we conclude that purely elastic and short range interactions lead to a universal power law scaling behaviour of the cell front roughness. The observed high values of the roughness exponent are in line with those seen for known universality classes of disordered elastic systems with short range interactions, such as one-dimensional interfaces subjected to thermal fluctuations (*ζ_th_* = 0.5) or random bond (*ζ_RB_* = 2/3) disorder [6, 31]. *In-vitro*, the sub-cell roughness scaling regime, with *ζ*_1_ ~ 0.58 is also coherent with such a description. However, the simulations fail to reproduce the lower roughness exponent value observed experimentally at few-cell lengthscales. This failure reinforces our hypothesis that a hierarchy of interactions is necessary to fully capture the behaviour of proliferating cell fronts, and that mid-lengthscale interactions are particularly important.

More generally, our approach extends literature applying models initially derived for physical systems to biological ones, demonstrating that many aspects can be described in purely physical terms. Defining different scaling exponents for interfacial roughening at different lengthscales during normal cell growth, and linking the evolution of these exponents to specific cellular pathways, represents a novel approach to understand the origins of pathological growth in deleterious situations, including cancer and traumatic injury.

## IV. METHODS

### A. Cell preparation

Rat-1 fibroblasts were maintained in Dulbecco’s Modified Eagle Medium (DMEM) with high glucose, L-glutamine, and phenol red (*Sigma-Aldrich*, Darmstadt, Germany #D5796), supplemented with 1% penicillin/streptomycin (*Sigma-Aldrich*, Darmstadt, Germany, #P4333) and 10% fetal bovine serum (FBS) (*Gibco Thermo Fisher*, Reinach, Switzerland, #10270106). Rat-1 fibroblasts were cultured at 37°C in a humidified incubator with 5% *CO*_2_ and were passaged 1:6 when they reached 80% confluence after washing with PBS and detaching with trypsin 10x (*Sigma-Aldrich*, Darmstadt, Germany #15400054). At the final passage, the cells were seeded around a prepositioned 6×2 cm^2^ insert (*Ibidi*, Gräfelfing, Germany) and incubated to confluence. The insert was removed and the resulting front allowed to settle and set for 2 hours prior to the first imaging steps.

GFP fluorescence marking of the cytoplasm was realised through retroviral modification using V21, premade LV vector, (vesicular stomatitis virus (VSV) glycoprotein (G) pseudotyped) obtained from the Viral Vector Facility of the Neuroscience Center Zurich. Nuclei were fluorescently marked using 32.0 micromolar Hoecht stain (*Sigma Chemicals*, USA, H 33258) for three hours.

For pharmacological modulation, we used (i) colchicine (*Sigma-Aldrich*, Darmstadt, Germany, #C9754), 0.32 μM concentration (ii) cytochalasine-B (Lucerna Chem AG, Luzern, Switzerland, #AGS-C-1027-5MG), 0.50μM concentration, (iii) meclofenamic acid (same as i and ii), and (iv) forskolin (same as i and ii). Employed concentrations were established in separate titrations with each inhibitor to determine effects on cell mortality, growth and morphology, as detailed in S1.

### B. Optical imaging

Brightfield phase and fluorescence microscopy imaging was carried out using a *Leica* DM6000 microscope coupled with a 1 Mpxs *Hamamatsu* CCD, at 1 px = 0.8μm resolution. To obtain the full panorama of the proliferating front, a frame matrix of 3 × 80 images, each 800 × 800 μm^2^ was captured every 4 hours, using software-controlled motorised stage steps. While in most cases the fluorescence signal remained relatively stable, for some fronts significant contrast variation artifacts made extraction of the imaged panorama extremely difficult. Such fronts, for instance the MFA-treated 36 and 40 hour timepoint, were not included in the dataset for analysis.

### C. Numerical methods

Numerical simulations of proliferating cell fronts were carried out using the *epicell* package, developed at the University of Geneva [32], adapted to the modeling of mechanical properties of two-dimensional cell sheets [24, 25].

Simulations were performed with the *freeBoundary* application, on a 20 × 1000 grid of two-dimensional hexagonal cells. The model was modified to favour divisions occurring just behind the cell front, in line with our empirical observations.

The cell area elasticity parameter *K* was set to 2.256 × 10^9^*N*/*m*^3^ following previous studies on mechanical properties of cell sheets, and the preferred cell area *A*^0^ was set to 3.14 × 10^−10^*m*^2^, following the experimental observations on the average size of the Rat1 cell line. The normalized inter-cell adhesion, 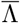, was explored in phase space from −1.975 to 0.5 in steps of 0.025. Likewise, the normalized cell perimeter contractility, 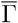, was explored in phase space from −0.3 to 0.675 in steps of 0.025.

4000 simulations were performed with a running time constraint of 12 hours and a maximum division number of 10^5^. Of these, 881 simulations were successful and were used as the basis for the discussion in the main text. Unsuccessful simulations are discussed in S8.

## V. ACKNOWLEDGEMENTS

This work was partially supported by the Commission Informatique of the University of Geneva. N.C. acknowledges support from the Federal Commission for Scholarships for Foreign Students for the Swiss Government Excellence Scholarship (ESKAS No. 2018.0636). The computations were performed at University of Geneva on the *Baobab* and *Mafalda* clusters. Lentivirus for GFP transfection was kindly provided by the Viral Vector Facility of the Neuroscience Center Zurich.

The authors declare no competing financial interests.

P.P. and S.A.B. designed and supervised the work. G.R performed the fluorescence measurements with assistance from A.R. and E.M. for cell culture and reagent preparation, and B.Z. for initial experiment implementation. I.G. carried out the numerical simulations. G.R., N.C., and I.G. analysed the data and wrote the manuscript with P.P. All authors contributed to the scientific discussion and manuscript revisions.

## SUPPORTING INFORMATION

Code used for the analysis is available in the form of Python scripts and Jupyter notebooks at https://gitlab.unige.ch/paruch-group/cell-fronts.

The data that support the findings of this study are openly available in Yareta at http://doi.org/10.26037/yareta:eyj33z7alje35cypydi5tecb2q in compliance with SNSF Data Management Plan guidelines.

### S1: Pharmacological modulator titration

To determine which concentration of the inhibitors would be most effective without significantly affecting cell mortality, separate titrations were conducted for each of the compounds on fully confluent healthy Rat1 cell cultures. Starting with values commonly used in our previous experiments, the applied concentration was increased in pseudo-logarithmic steps (1x, 2x, 5x, 10x, 20x, etc), up to a killing concentration determined by visual inspection of the cells after 24 hours of exposure, in comparison with cells prepared at the same time under identical conditions without inhibitor exposure.

Inhibitor effects on cell growth, morphology, and mortality were evaluated based on the presence and extension of lamelipodia, the distance between cell nuclei, loss of confluence, and of course the amount of dead cells. For colchicine, the initial concentration of 0.32 μM did not affect cell mortality, but morphology effects of a more rounded and globular cell shape were beginning to be apparent. At higher concentrations, these morphology changes were more significant, but we also observed significant mortality increase above 0.625 μM, leading us to conclude that the original concentration of 0.316 μM was already at the optimal value. For the cytochalasin B, starting from a concentration of 0.1 μM, cell mortality increased sharply from less than 5% to more than 30% for 0.5 to 1 μM, and the colony was seriously depopulated (50% mortality at 2 μM). We therefore chose an effective concentration of 0.5 μM. The toxicity of meclofenamic acid was likewise easily decided, the mortality again increasing rapidly from less than 5% to more than 80% between 50 and 100 μM, leading us to chose an effective concentration of 50 μM. Finally, with forskolin, we did not observe significant effects on cell mortality. At 5 μM we began to see apparent elongation of the cells. These effects appeared to saturate at 10 μM, which was therefore the concentration chosen for the experiments. We generally observed that at concentrations approaching the killing concentration, cells appear to shrink and we begin to observe loss of confluence with increasing amounts of empty space between the cells. The cells also detached readily from the substrate surface, dying and floating above the rest of the colony.

### S2: Image processing

Each of the 1000 × 1000 px images acquired using the Leica DM6000 were indexed by the *x, y* position of the motorised sample stage (the vertical height *z* was kept constant once the desired focal plane was established), allowing us to stitch the individual images in the frame matrix into a full panorama of the proliferating front, as shown in Fig. S1(a), using an in-house developed Python algorithm. To fully discriminate the cell colony from the background the original greyscale fluorescence images are binarised using the Otsu algorithm and the brightness histogram for the panorama, as seen in Fig. S1(b), with the colony in white. As a result of varying GFP expression between cells, the binarisation still leaves some dark areas behind the cell front, necessitating a “hole filling” step carried out with the Scipy function, “binary_fill_holes”, which leads to the image shown in Fig. S1(c). In this image, we identify the cell culture as the largest group of connected white pixels, obtained using the Scikit “label” function. The set of *x*, *y* coordinates of the front position is finally obtained using the Scikit function “find_contours”, and can be seen in Fig. S1(d), overlaid on the original fluorescence microscopy image the cell colony.

**FIG. S1.**
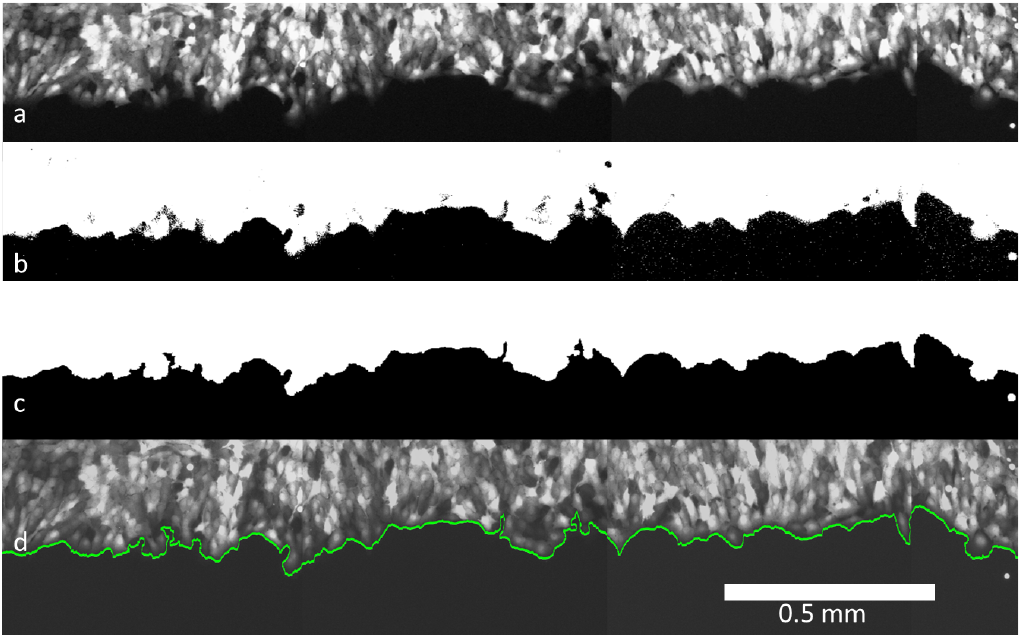
Image processing to obtain front position. The fluorescence microscopy image (a) is binarised using the Otsu algorithm (b), and fluorescence variations corrected with a “hole filling” step (c). The colony is identified as the largest connected island on the image and the front position (green) is extracted as its lower contour, shown overlaid on the original fluorescence image (d).

**TABLE S1.** Results of individual toxicity titrations to establish the final concentration (indicated in bold) of the inhibitors used in the study.

|  |  |
| --- | --- |
| Forskolin | Cell mortality |
| 2 $\mu\text{M}$ | Negligible (<5%) |
| 5 $\mu\text{M}$ | Negligible (<5%) |
| **10 $\mu\text{M}$** | Negligible (<5%) |
| 20 $\mu\text{M}$ | Significant (30%) |
| Colchicine | Cell mortality |
| **0.32 $\mu\text{M}$** | Negligible (<5%) |
| 0.63 $\mu\text{M}$ | Significant (10%) |
| 1.25 $\mu\text{M}$ | Significant (50%) |
| 3.20 $\mu\text{M}$ | Significant (90%) |
| Cytochalasin B | Cell mortality |
| 0.1 $\mu\text{M}$ | Negligible (<5%) |
| 0.25 $\mu\text{M}$ | Negligible (<5%) |
| **0.50 $\mu\text{M}$** | Negligible (<5%) |
| 1.00 $\mu\text{M}$ | Significant (30%) |
| 2.00 $\mu\text{M}$ | Significant (50%) |
| MFA | Cell mortality |
| 10 $\mu\text{M}$ | Negligible (<5%) |
| 25 $\mu\text{M}$ | Negligible (<5%) |
| **50 $\mu\text{M}$** | Negligible (<5%) |
| 100 $\mu\text{M}$ | Significant (80%) |
| 250 $\mu\text{M}$ | Significant (80+%) |

### S3: Monovalued approximation

The extracted front position follows the contours of the individual cells, and shows small overhang features where a single cell or small clusters of a few cells at the edge of the colony orient somewhat diagonally. However, the framework of disordered elastic systems, with an analysis of the correlation function *B*(*r*) of relative displacements Δ*u*(*r*), is only applicable in the case of a univalued interface, meaning that elements such as overhangs, holes, and islands cannot be treated within this approach. We therefore approximate the extracted front position by a univalued function. This correction is carried out using 3 different approaches, as shown in Fig. S2: ‘outer’ contour (yellow), choosing the highest value of *u*(*z*) if multiple points of the front are present at a given lateral coordinate *z*; ‘inner’ contour (green), choosing the lowest value of *u*(*z*) if multiple points of the front are present at a given lateral coordinate *z*; and ‘sum’ contour (blue), adding all the pixels of the colony at a given lateral coordinate *z* to obtain an effective univalued *u*(*r*).

As can be seen in Fig. S2(a), the roughness *B*(*r*) calculated for each of these corrected fronts shows small differences at sub-cell lengthscales, but these become insignificant by the time the behaviour at few-cell lengthscales is examined. Since the sum approach effectively smooths the interface, to preserve at least partially the effect of the overhangs, which are clearly a feature of the biological system, we chose to use the outer contour for all the analysis presented in this study.

This results in abrupt but still univalued ‘jumps’ in the front position. In order to estimate the impact of these jumps on the front geometry, we subtracted the outer and inner contours of the front, and extracted the statistics of the resulting deviations from a straight line. As can be seen in Fig. S3(a,b), the jumps, reflecting the original overhangs, are generally limited to at most a width of 8 μm (i.e. less than the characteristic cell size) and an average height of 200 μm (i.e. approximately 10-12 cells).

**FIG. S2.**
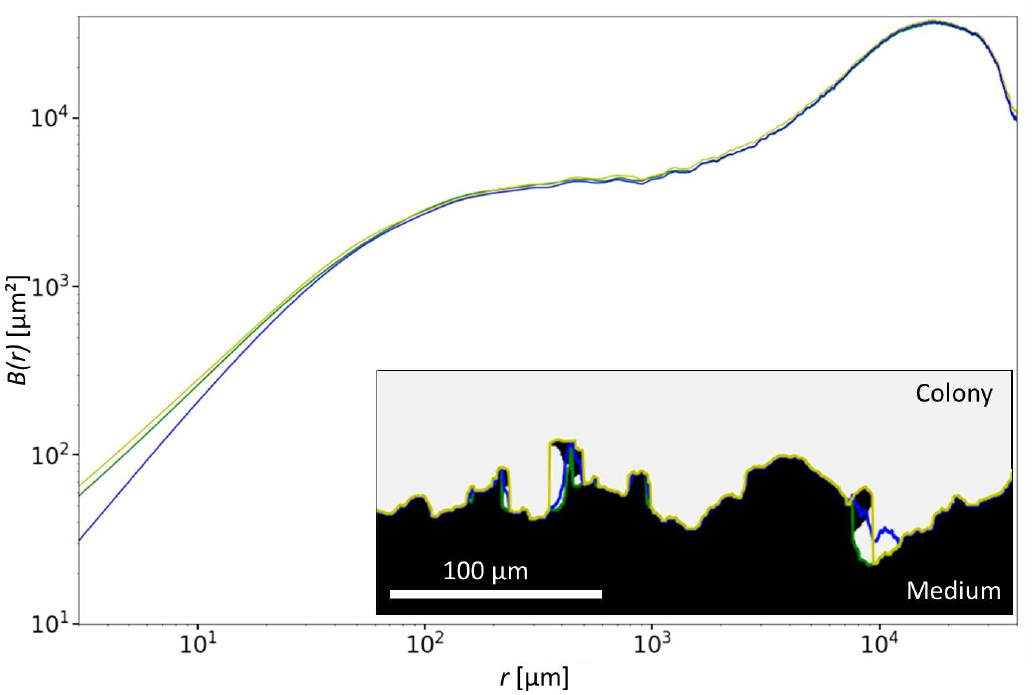
Univalued correction of front position. (a) Cell front roughness *B*(*r*) extracted after correction of the intermittent overhangs using the ‘outer’ (yellow), ‘inner’ (green), and ‘sum’ (blue) contours shows only very minor differences at subcell lengthscales.

**FIG. S3.**
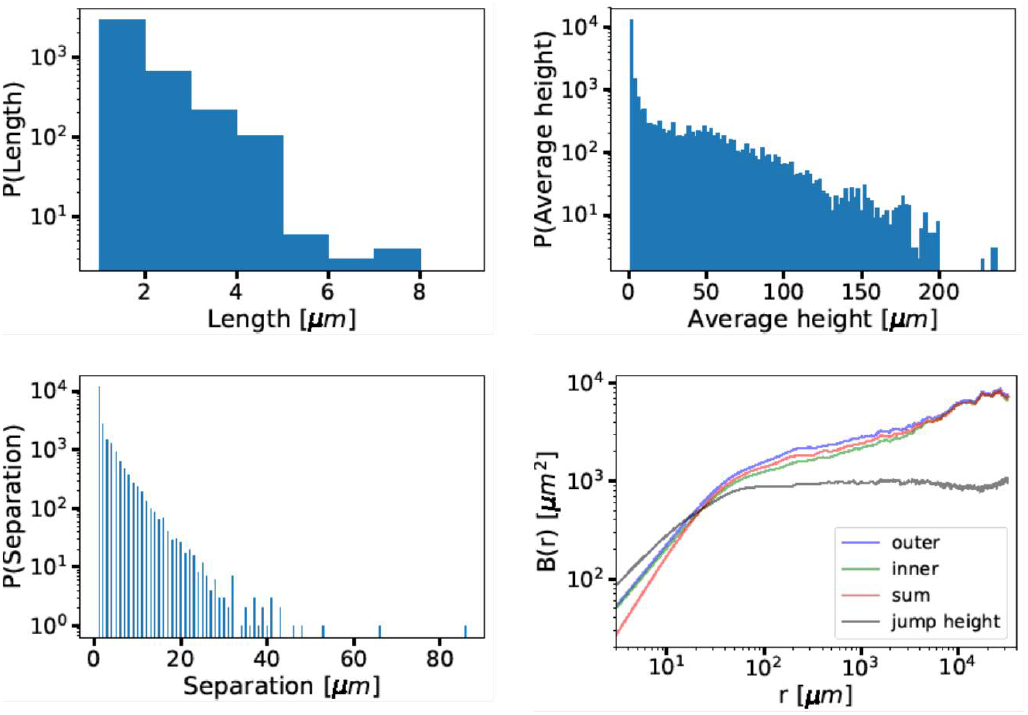
Statistics of jump features resulting from the correction of overhangs in the cell front position. (a) Distribution of jump widths along the cell front. No overhangs wider than approximately a half cell size are observed. (b) Distribution of average jump heights perpendicular to the cell front. (c) Distribution of the interjump separations along the cell front.

### S4: Rounding of the initial configuration

During the lift-off patterning, the flexibility of the silicone insert used to demarcate the initial position of the cell front in a confluent cell culture can lead to a rounded arc geometry rather than a straight line configuration. This arc corresponds to a segment of a circle with a radius of approximately 1 m, which can be quite precisely overlaid over the initial front position, as shown in Fig. S4(a), and contributes an identifiable artefact to the calculation of the front roughness *B*(*r*). As can be seen in Fig. S4, *B*(*r*) for such a 1 m radius circle arc shows a power law growth, with a roughness exponent *ζ* = 1. However, the initial roughness levels are over 4 orders of magnitude lower than that of the cell front itself, starting at approximately 2 × 10^−4^ μm^2^ at *r* = 1 μm. The roughness increases to 3 × 10^4^ at *r* = 30mm, the highest available lengthscale of the analysis. While this artefact, when present, becomes the dominant contribution to the experimental observations above *r* = 2 mm, and affects to a lesser degree the roughness of the cell front above 500 μm, it does not affect our analysis of the double power law growth behaviour with very different roughness exponents at subcell (4-18 μm) and few-cell lengthscales (80-180 μm).

**FIG. S4.**
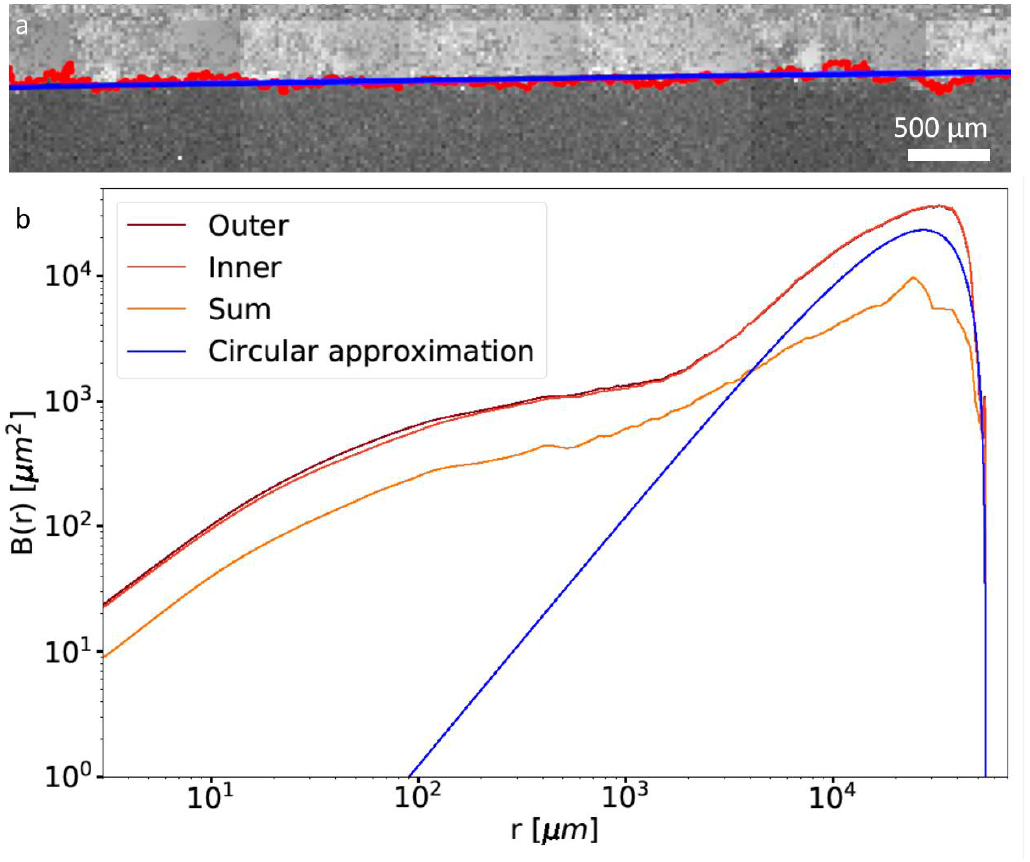
(a) Fluorescence images of the cell front (GFP-stained cytoplasm), with the extracted front position (red), overlaid with the circle segment approximation (blue). (b) Roughness *B*(*r*) calculated for the three different overhang correction approaches, and of the circle segment used to approximate the initial position resulting from a slight bending of the flexible s ilicone insert.

### S5: Reproducibility of roughness analysis

Our previous studies of physical interfaces have shown that sufficiently long individual fronts, or sufficient different iterations of shorter fronts are necessary to accurately extract the value of the roughness exponent *ζ* [33, 34]. The relatively large cell front panoramas obtained in our experiments (length *L* ~ 2^16^ px or ~ 5cm) provide ample statistics for the analysis of the roughness *B*(*r*), restricted to 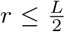, and thus a high level of confidence in its accuracy.

As a further test, and to check the reproducibility of the experimental protocol, we repeated each experiment three times under control conditions to obtain the time evolution of three separate cell front panoramas over 40 hours. For each of these realizations *i* (*i* = 1, 2, 3), we obtain the functions *u_i,t_*(*r*) describing the position of the cell-front after an evolution time *t* = 0, 4, 8, ⋯ 40 hours. We compute the roughness *B*_*i,t*_(*r*) for each of these fronts, and the average 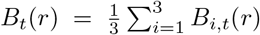. over the three realizations *i* at the same time *t*. As shown in Fig. S4, the roughness does not appear to change significantly from one realisation to the other (apart from the rounding artefact present in one of the realisations, as discussed in Sect. S4), and after a long evolution time, the roughness obtained for the three realisations appears to converge to the average value.

### S6: Extracting the roughness exponent

One of the most important features of the roughness function *B*(*r*) is that it provides information about the interactions present in a system. To capture this information, two correlated elements are necessary. First, the spatial regions where the function follows a power-law need to be identified, and second, the fit of the function with a power-law should be done over this region to extract the roughness exponent describing the interactions at this length-scale.

In the theory of disordered elastic systems, where, for example, a system evolves as result of the competition between the elasticity of the interface *c*, the disorder or pinning, and thermal fluctuations proportional to the temperature *T*, it is well known that the roughness function at short length-scales is dominated by thermal fluctuations, and follows a power law behavior 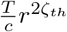, where the roughness exponent is given by the thermal exponent *ζ*_*th*_ = 0.5. However, at large length-scales *B*(*r*) is dominated by disorder, if present, and depending on the disorder type follows a different power-law proportional to 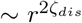. For example, for random bond disorder the scaling is governed by 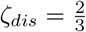.

In the case of biological systems, the nature of the interactions between cells and how they affect the dynamics and statics of a cell front is not yet well established, and is what motivates the current study. In order to apply the theory of disordered elastic systems to effectively capture the information about the interactions in a growing colony of cells, we identify the regions where *B*(*r*) follows a power-law behavior as follows: we perform a weighted fit of log *B*(*r*) with a linear function of *r* over the region [log *r_i_*, log(*r_i_* + *w_i_*)], where *r_i_* is the starting point and *w_i_* is the length of the fitting region chosen according to the initial value *r_i_*, in order to always obtain a window of equivalent size on a logarithmic scale for each starting point (see Fig. S6). The weights used for the fit are computed as the uncertainties Δ*y* associated to each point of *y* = log *B*(*r*). These uncertainties can be estimated as 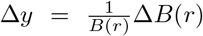, where 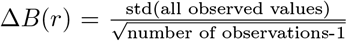, and the number of observations decreases when *r* is increased (observations=total interface length − *r*). For each fit we compute the goodness of the fit through *R*^2^.

As can be seen in Fig. S7, which shows *R*^2^ vales for fits of the roughness *B*(*r*) at *t* = 0 hours averaged for the three fronts under control conditions, two regions emerge where the power law fits with *B*(*r*) ~ *r*^*ζ*^ are highly pertinent. At short lengthscales (for small *r*_*i*_) we obtain very high *R*^2^ values above 0.9998, even for relatively large window sizes *w*_*i*_. *R*^2^ then abruptly decreases, and a second local maximum with values around 0.9992 is reached at ~ 80μm. This behaviour suggests two separate power law scaling regimes are present in *B*(*r*).

To chose the best window size for each region we complemented the mathematical consideration of *R*^2^ with additional biological arguments. For *r_i_* = 4μm, even if the goodness of the fit is high up to 50μm, we restricted the fit to not exceed the average length of a cell ~20 μm, in order to capture the behavior of *B*(*r*) at subcell lengthscales, thus defining a first region *I* = [4, 16]μ m. For the second region, we considered that important mechanisms of intercell communication, in a wide range of mammalian epithelial cells, have a range of at most a few cell lengths for example the cell networks reported for gap junction communication in rat kidney cells[29] as well as mechanical interactions via the cytoskeleton and force transfer via the substrate, which were found to decrease linearly with distance [14]. We note here that the inhomogeneous distribution of cell division we observed, which seemed to be more prevalent in cells close, but not directly at the colony edge, can also be considered as an interaction at this range. We therefore restricted the second region to *II* = [80, 180]μm. For all the experimental cell fronts, fits of *B*(*r*) were performed over these two regions *I* and *II* to obtain *ζ_I_* and *ζ_II_*.

### S7: Effective flattening by filling of cavities

During the measurements, we occasionally found large cavities at the edge of the colony, as shown in Fig. S8(a). The presence of these features significantly affected the dynamics of the colony, with cells rapidly moving towards the cavity and progressively filling it, as can be seen in Fig. S8(b,c), with the inflowing cells aligned radially towards the cavity center. Only when the cavity was filled did this section of the front continue to move outward again, as previously observed by Kim *et. al.* [35]. While we excluded affected segments of the fronts containing such features from the statistical analysis of roughness and dynamics, we believe that they are potentially related to the steady-state saturation of *B*(*r*) at higher lengthscales.

**FIG. S5.**
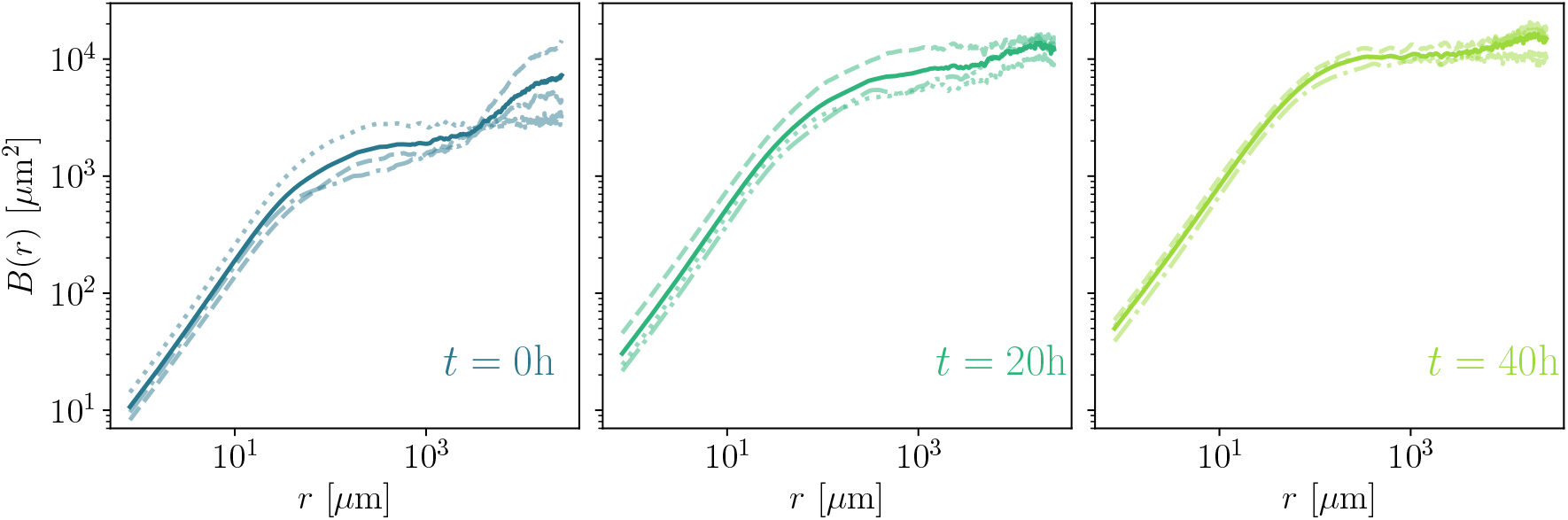
Reproducibility of roughness analysis. Roughness function *B*(*r*) computed for three experimental realisations of the cell front under control conditions (discontinuous lines), and the average roughness (solid line) at different evolution times *t* = 0, 20, 40 hours.

**FIG. S6.**
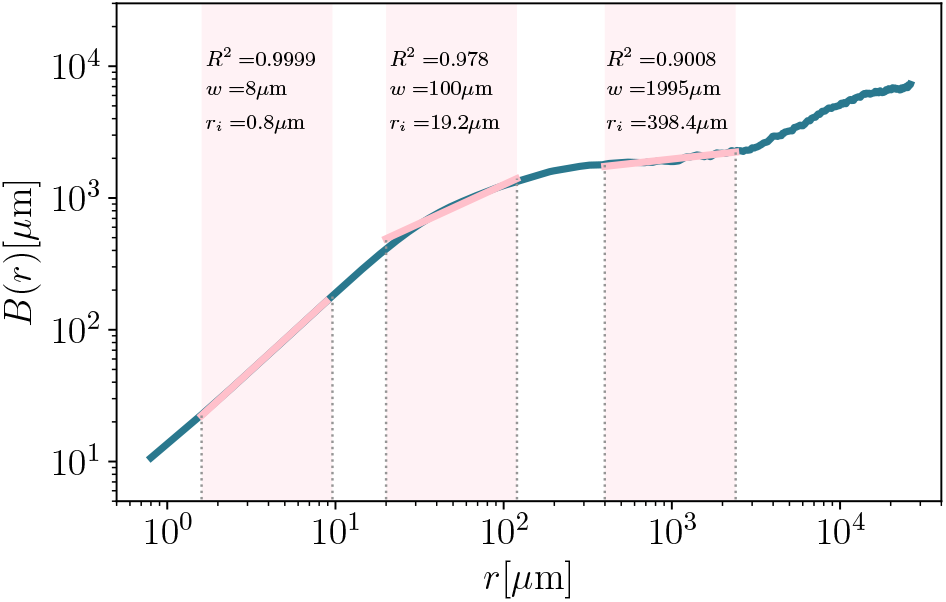
To determine the regions where the roughness function follows a power-law behavior, we explored different ranges [*r*_*i*_, *r*_*i*_ + *w*_*i*_] to fit *B*(*r*). Here, 3 different power-law fits of the roughness of the initial front configuration at *t* = 0 under control conditions are shown, for *r*_*i*_ = 0.8, 19.2, and 398.4μ*m*, and equivalent window-sizes on a logarithmic scale *w*_*i*_ = 8, 100, and 1995μ*m*. We measured the goodness of the fit through *R*^2^.

**FIG. S7.**
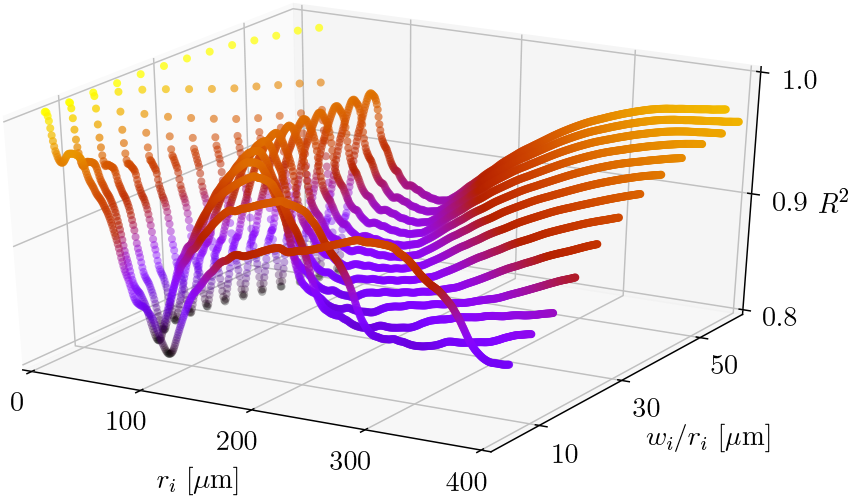
Identifying power law scaling regimes. *R*^2^ obtained for the weighted fit of log *B*(*r*) averaged for the three fronts under control conditions at *t* = 0. The fit was performed over the regions [*r*_*i*_, *r*_*i*_ + *w*_*i*_], where *w*_*i*_ was chosen for each *r*_*i*_ to obtain an equivalent window size on a logarithmic scale, independently of *r*_*i*_. Two regions with high *R*^2^ are observed, one at short length-scales, and the other with a local maximum at *r*_*i*_ ~ 80μm, separated by a region of low *r*^2^, suggesting two separate power law scaling regimes.

### S8: Numerical simulations

The numerical simulations of proliferating cell fronts were performed by using the *epicell* package, developed at the University of Geneva [32], adapted to the modeling of mechanical properties of two-dimensional cell sheets[24, 25].

The *epicell* model used is based on solving the following dimensionless Hamiltonian on a regular grid of two-dimensional hexagonal cells:

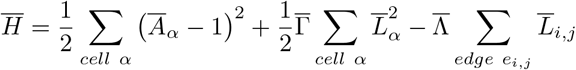

with 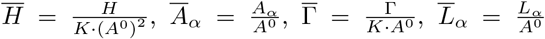, 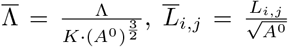, *H* the Hamiltonian, *A*^0^ the preferred cell area, *K* the cell area elasticity, Γ the cell perimeter contractility, Λ the inter-cell adhesion represented by the cell-cell edge tension, *A_α_* the area of cell *α*, *L_i,j_* the length of the inter-cell edge *e_i,j_*. The renormalization of the Hamiltonian with the constraint that all cells are of the same fixed type, elasticity and preferred size, yields a two-variable system, with reduced parameters 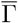 and 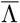.

Additionally, the model was modified to include a nontrivial cell division descriptor, favoring divisions to occur just behind the cell front, in line with out empirical observations. This was implemented through setting a weight prefactor to the division probability following *W*_*div*_ = 0 at the cell front (*d* = 0) and 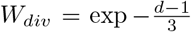 for cells with distance *d* behind the front.

**FIG. S8.**
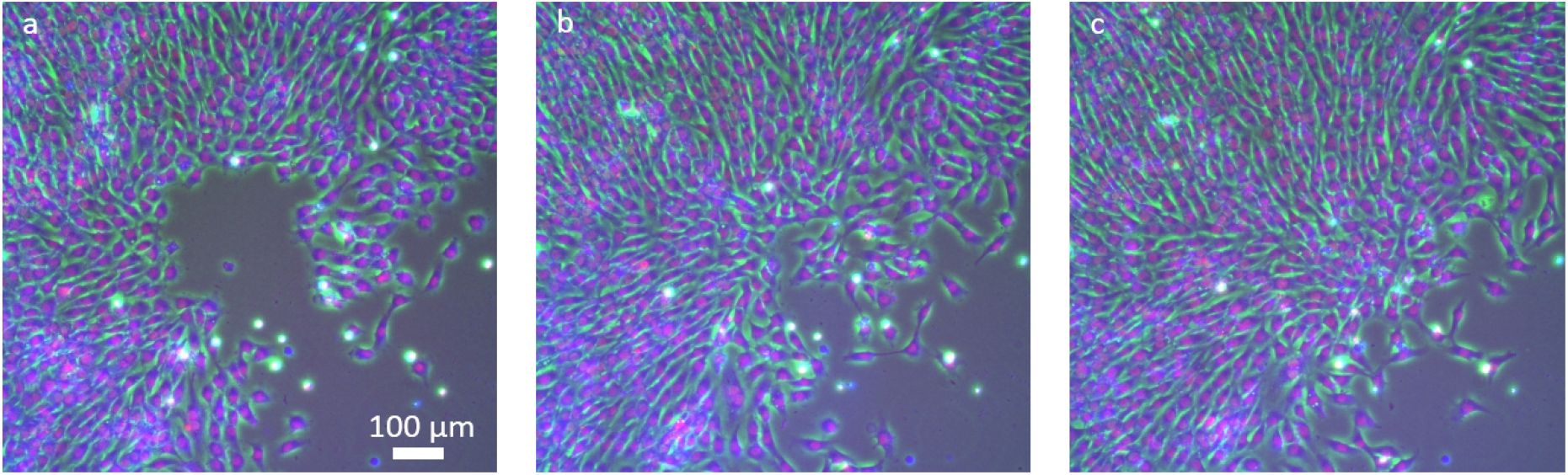
Cavity filling by collective/directed cell flow. Fluorescence images of a cell front, imaged every 4 hours, with a cavity initially present (a), progressively filled by the rapid inflow of cells from the bulk of the colony (b). Only upon complete filling of the cavity (c) does the front start to move outward again.

#### Parameters and execution

Seventeen parameters are necessary to run the *free-Boundary* application included in the *epicell* package. They are listed below with their description and values taken for the current study:

- **modelId**: The model dictates which cells are selected for division in priority. Here, set to 3, so that the largest and less compressed cells would divide first.
- **nRep**: Number of replicates of the cell front, here set to 1.
- **myseed**: Seed for the random number generator, here set to 1.
- **K**: The cell area elasticity parameter *K*. Set to 2.256 · 10^9^*N/m*^3^ following previous studies on mechanical properties of cell sheets.
- **A0**: Preferred cell area *A*^0^. Set to 314 · 10^−10^*m*^2^, following the experimental observations on the average size of the Rat1 cell line.
- **lambdaNorm**: Normalized inter-cell adhesion, 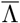. Parameter explored in phase space from −1.975 to 0.5 in steps of 0.025.
- **gammaNorm**: Normalized cell perimeter contractility, 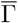. Parameter explored in phase space from −0.3 to 0.675 in steps of 0.025.
- **ncols**: Number of columns of cells within the cell sheet. Set to 20 here to have a thin but long cell sheet.
- **nrows**: Number of rows of cells within the cell sheet. Set to 10^3^ here to have a thin but long cell sheet.
- **xmin**: Left hard boundary position. Set to 0 here so that the system imposes boundaries automatically after the initial relaxation step.
- **xmax**: Right hard boundary position. Set to 0 here so that the system imposes boundaries automatically after the initial relaxation step.
- **ymin**: Bottom hard boundary position. Set to 0 here so that the system imposes boundaries automatically after the initial relaxation step.
- **nbDivMax**: The maximum number of cell divisions before the simulation terminates. Set here to 10^5^ to allow a large number of divisions compared to the number of cells in the sheet.
- **elasticCoef**: Elastic coefficient of the hard boundaries, used in this work to mimic an infinite incompressible cell sheet on three of the sides as to promote growth along one direction. Set to 10 here to provide a hard boundary.
- **sdLambda**: Standard deviation of the inter-cel following the experimental observations on the average size of the Rat1 cell line.
- **sdGamma**: Standard deviation of the cell perimeter contractility Γ. Set to 0 in the current study.
- **boundaryLambda**: Normalized line tension of the cell sheet edges, used for simulating an interaction of the cells with the external media. Set to 0 in the current study.

The *freeBoundary* code was executed on the University of Geneva cluster, *Baobab*, with a time constraint before termination of 12 hours. The 4000 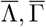 parameter pairs were computed in parallel and 881 simulations were successful. Unsuccessful simulations were either due to nonphysical parameters (1411 simulations), impossible convergence at the initial relaxation stage (1050 simulations), nonphysical mechanical behaviour of vertices (651 simulations), and finally unexpected simulation terminations due to specific pathological parameter sets (7 simulations). The outcome of all parameter pairs is shown in Fig. S9, with successful simulations labeled as *OK*.

**FIG. S9.**
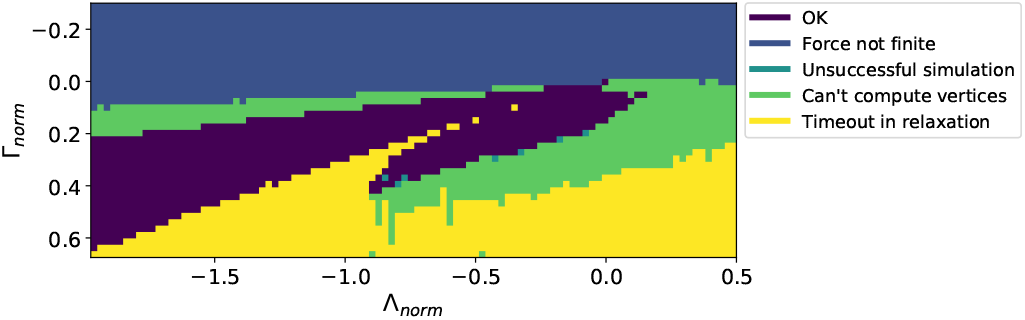
Complete simulation space with 4000 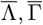 parameter pairs. Out of this, 881 simulations were successful, labeled as *OK*. The unsuccessful simulations either occurred due to nonphysical parameters (1411 with infinite forces and 651 with nonphysical vertices) or due to non-convergence during the initial relaxation step (1050 simulations). A minority, 7 simulations, ended unexpectedly due to specific pathological parameter sets.

#### Phase diagram investigation

Our study focused on the successfully simulated region of the phase diagram, as shown in Fig. S10. The phase diagrams for the asymptotic roughness exponent *ζ*, quality of fit *R*^2^ for the asymptotic exponent, total front displacement from its initial position, velocity of the front over the final 100000 iterations, and the number of iterations for each simulation are shown in Fig. S10(a,c,e,g,i) respectively.

As discussed in the main text, the analysis of simulation parameters enables the extraction of two branches of physical observables, with the lower branch exhibiting pathological behaviour including quasi-nonexistent front displacement/velocity, periodic features in the roughness, and large error during extraction of the roughness exponent *ζ*. Through the combination of the phase diagrams, a region of *stability* was defined visually as a red bounding polygon, with edges located at 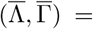 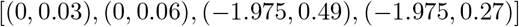. This filtering enables the removal of pathological scenarios, and allows for a better selection of the appropriate mod-els for comparison with *invitro* experiments. For instance, the overall average roughness exponent 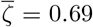 as shown in the distribution Fig. S10(b) is highly asymmetric, whereas sampling *ζ* values in the region of stability yields an average roughness exponent 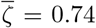 with a smaller error and a more symmetric distribution.

It is also noteworthy that whilst *ζ* and the number of iterations following the initial relaxation step (Fig. S10(i,j)) seem to be concentrated around specific values, the total displacement and final velocity in Fig. S10(e,f,g,h) are more evenly distributed, and in fact tend to increase with decreasing inter-cell adhesion and cell contractility.

**FIG. S10.**
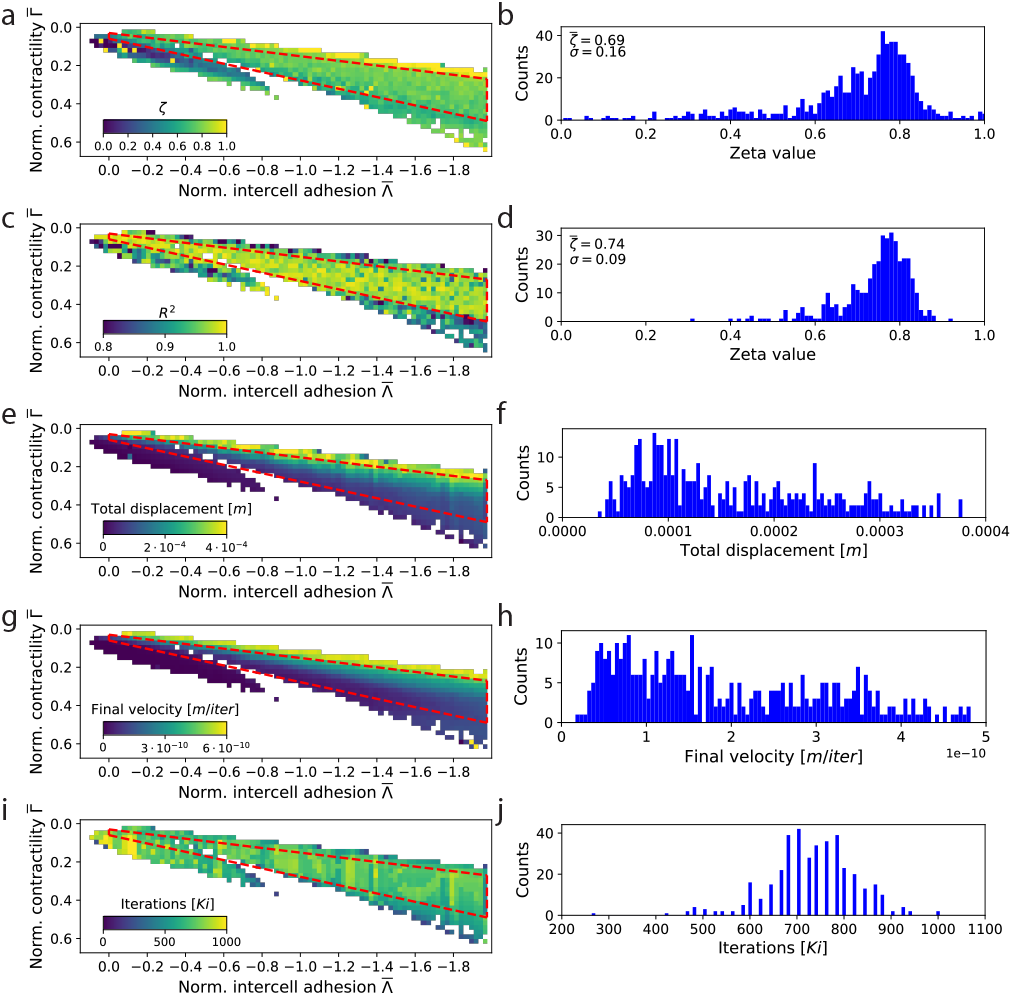
Simulation observable phase diagrams and their corresponding distributions. The red polygons delimit the region of stability defined from visual observation of the extracted physical parameters. (a) Phase diagram of the asymptotic roughness exponent *ζ*, computed by fitting the time evolution of the roughness exponents of fronts. (b) Distribution of the *ζ* values, showing a large spread, asymmetry and a mean at 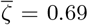. (c) Goodness-of-fit estimate *R*^2^ for the *ζ* fits, demonstrating a large region of high quality fits within the region of stability. (d) Distribution of *ζ* values within the region of stability, with a mean of 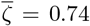. (e) Phase diagram and (f) region of stability distribution of the total front displacement. (g) Phase diagram and (h) region of stability distribution of the final front velocity. (i) Phase diagram and region of stability distribution of the number of iterations for each simulation parameter pair following initial relaxation.

## Notes

### Competing Interest Statement

The authors have declared no competing interest.

